# Preservation of thalamocortical circuitry is essential for good recovery in comatose survivors of cardiac arrest

**DOI:** 10.1101/2022.11.02.514844

**Authors:** Prejaas K. Tewarie, Marleen C. Tjepkema-Cloostermans, Romesh G. Abeysuriya, Jeannette Hofmeijer, Michel J.A.M. van Putten

## Abstract

Continuous EEG monitoring contributes to prediction of neurological outcome in comatose cardiac arrest survivors. While the phenomenology of EEG abnormalities in postanoxic encephalopathy is well-known, the pathophysiology, especially the presumed role of selective synaptic failure is less understood. To further this understanding, we estimate biophysical model parameters from the EEG power spectra from individual patients with a good or poor recovery from a postanoxic encephalopathy. This biophysical model includes intracortical, intrathalamic, and corticothalamic synaptic strengths, as well as synaptic time constants and axonal conduction delays. We used continuous EEG measurements from hundred comatose patients recorded during the first 48 hours post-cardiac arrest, fifty with a poor neurological outcome (Cerebral Performance Category (CPC=5)) and fifty with a good neurological outcome (CPC=1). We only included patients that developed (dis-) continuous EEG activity within 48 hours post-cardiac arrest. For patients with a good outcome, we observed an initial relative excitation in the corticothalamic loop and corticothalamic propagation that subsequently evolved towards values observed in healthy controls. For patients with a poor outcome, we observed an initial increase in the cortical excitation-inhibition ratio, increased relative inhibition in the corticothalamic loop, delayed corticothalamic propagation of neuronal activity, and severely prolonged synaptic time constants, that did not return to physiological values. We conclude that the abnormal EEG evolution in patients with a poor neurological recovery after cardiac arrest may result from persistent and selective synaptic failure that includes corticothalamic circuitry, but also delayed corticothalamic propagation.

## Introduction

Continuous EEG monitoring in the first 24 hours after cardiac arrest has a pivotal role in prognostication in comatose survivors after cardiac arrest^1^, allowing reliable prediction of neurological outcome in about 50% of the patients^2^. Neurological prognostication not only depends on identification of specific EEG abnormalities, but also on the evolution and timing of these abnormalities post-cardiac arrest^3^. In all comatose patients directly after cardiac arrest the EEG is severely disturbed and mostly suppressed. In patients with a good neurological outcome, EEG activity typically evolves towards continuous brain activity on time scales of 24 hours^4, 5^. A delayed (*>* 24 h) evolution towards continuous activity is associated with a poorer outcome^1^. Persistent suppression with or without synchronous bursts, or generalized periodic discharges (GPDs) on a flat background are invariably associated with a poor outcome^2, 6, 7^.

While the clinical relevance of EEG phenomenology in postanoxic encephalopathy is well-known, the underlying patho-physiological mechanisms of EEG patterns associated with poor or good outcome are incompletely understood. Only in extreme cases, the pathophysiology is relatively clear: if ATP is depleted sufficiently long, irreversible loss of resting membrane potentials occurs, and cell swelling will follow, resulting in massive neuronal death accompanied by persistent iso-electric EEG^8–12^. However, in the pathogenesis of rhythmic and periodic or diffusely slowed EEG, as observed in many patients with moderate to severe hypoxic/ischaemic injury, it is predominantly synaptic failure that results in these abnormal EEG patterns^12–14^, where our understanding of selective dysfunction of excitatory or inhibitory synapses is incomplete^6^.

Only a few studies have explored this role of synaptic damage in EEG abnormalities in postanoxic encephalopathy. These studies were mainly based on so-called mean-field models^15, 16^. Several of these models can describe average post-synaptic membrane potentials of coupled excitatory and inhibitory cortical populations as a function of spike rates, synaptic strengths and synaptic time constants. The voltage fluctuations of the excitatory cortical neurons can subsequently be used as a proxy for EEG registrations. A common approach for mean-field modelling is hypothesis driven manipulations of model parameters^14, 17, 18^. For instance, selective elimination of excitatory synapses from pyramidal neurons to inhibitory neurons in a mean-field model showed GPDs that resembled empirically observed GPDs^17^. Another modelling study on postanoxic encephalopathy showed that a gradual recovery of synaptic strength led to an EEG evolution that was consistent with empirical data, i.e. suppressed activity, followed by burst-suppression and continuous activity. Here, the amount of suppression was a function of the “hypoxic burden” (depth and duration). This “hypoxic burden” determined whether continuous activity could be retrieved as final stage^14^.

Though very informative in providing a qualitative explanation of the evolution of EEG abnormalities, these previous approaches cannot retrieve subject specific trajectories of model parameters, such as temporal changes in synaptic properties. This contrasts with a recent approach using a corticothalamic mean-field model where model parameters are retrieved by finding the best fit between the observed and model’s power spectrum for every point in time, enabling real-time tracking of mean-field parameters over time for each individual patient^19^. The model has successfully been used to track sleep stages in terms of temporal evolution of physiological parameters^19^. The use of this parameter estimation method yields time-resolved trajectories of both excitatory and inhibitory synaptic strengths and synaptic time constants. This method also estimates non-synaptic parameters such as corticothalamic conduction delays. It therefore allows us to test the hypothesis that EEG abnormalities in patients after cardiac arrest can be completely explained by isolated selective synaptic failure. Here, we use this method to study pathophysiological mechanisms of EEG pattern evolution in comatose patients with a postanoxic encephalopathy.

## Results

We used EEG data from 100 healthy control subjects to obtain reference values for all estimated model parameters. We selected EEGs from 50 comatose survivors of cardiac arrest with a poor outcome and 50 patients with a good outcome from our previously published data-set^3, 6^. We only selected patients who developed a discontinuous or continuous EEG within 48 hours after cardiac arrest (see methods section for rationale). Figure 1A shows an example of EEG segments of 15 seconds from two patients at t=12 and t=24 hours after cardiac arrest as example. The EEG in the upper panel shows an evolution from burst-suppression at 12 hours after cardiac arrest to a continuous EEG with alpha activity (8-13 Hz) at 24 hours after cardiac arrest from a patient with a good outcome. The EEG in the lower panel shows an evolution from burst-suppression at 12 hours after cardiac arrest to a nearly continuous EEG with slower theta rhythms (4-8 Hz) at 24 hours after cardiac arrest from a patient with a poor outcome. For every subject, we derived a power spectrum for every hour of their EEG recording until 48 hours post-cardiac arrest. Figure 1B shows the evolution of the power spectrum for the same patients as in Figure 1A with a good (upper panel) and a poor outcome (lower panel). The occurrence and a shift of the spectral peak in the alpha band is clearly visible in the patient with a good outcome (purple corresponds to t=12 h and light green to t=48 h). A similar evolution is visible in the patient with poor outcome, but the spectral peak remains in the theta band.

**Figure 1.**
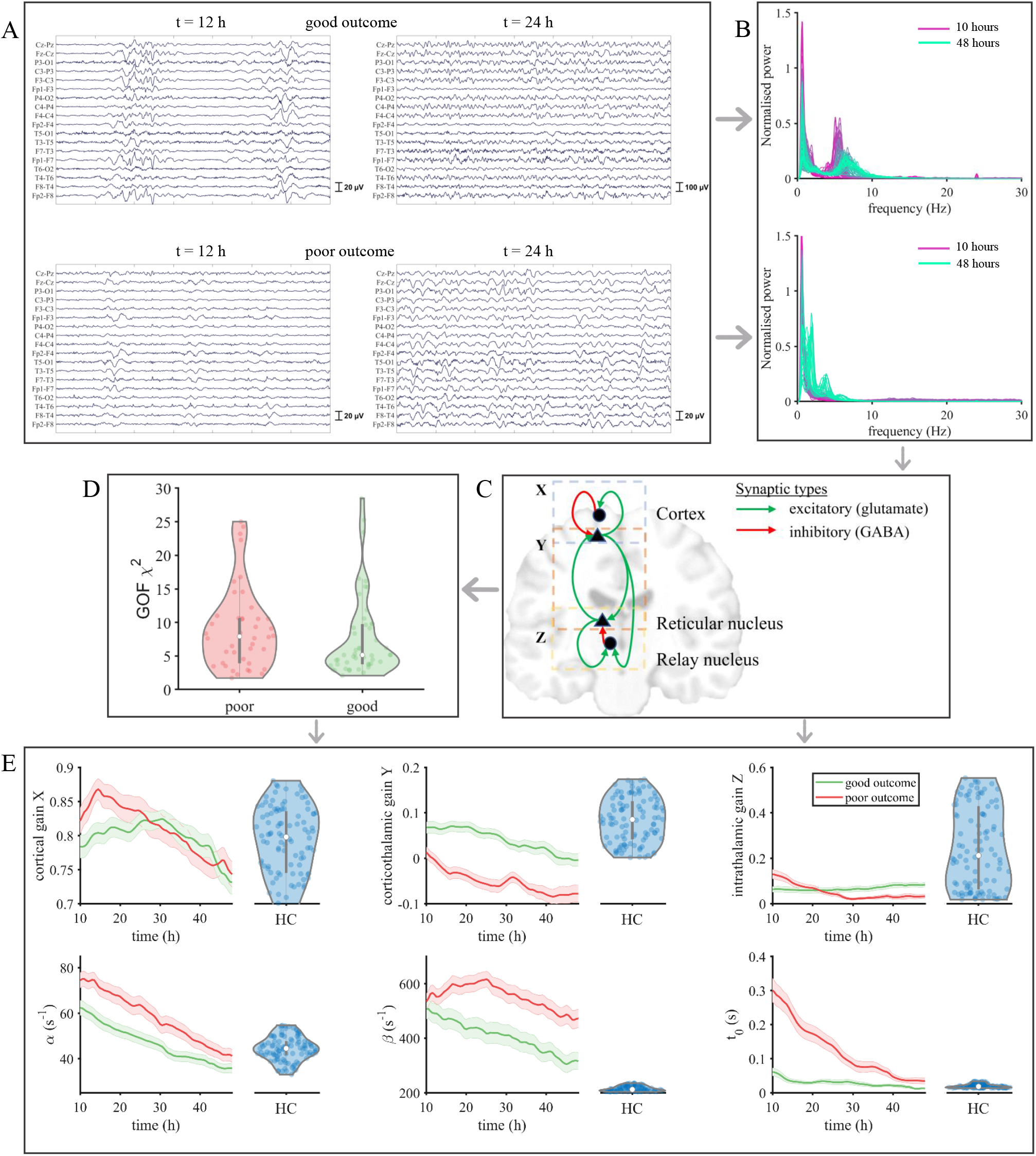
Estimating model parameters from EEG data. Panel A shows two EEG segments from two subjects at t=12 hours and t=24 hours after cardiac arrest. Upper panel shows EEG data from one subject with good outcome, lower panel shows EEG data from a different subject with poor outcome. The consecutive power spectra for the same subjects are demonstrated in panel B. Power spectra from every hour are depicted, starting at 10 hours after cardiac arrest (purple) to 48 hours after cardiac arrest (light green). Panel C shows the corticothalamic mean-field model. There are two populations in the thalamus and cortex. Red lines correspond to inhibitory synaptic connections and green lines to excitatory synaptic connections. Panel D shows the distributions (violin plots) of the goodness-of-fit (GOF) *χ*^2^ for the two groups. Here, every dot corresponds to the mean GOF across time points for one subject. Panel E shows the mean and standard error for all parameters, alongside the distribution of values from healthy controls (blue violin plots). Key findings in patients with a poor outcome are 1) an initial high cortical excitation inhibition ratio; 2) a persistent loss of corticothalamic gain *Y*, i.e. relative excess of corticothalamic inhibition; 2) slow synaptic responses (high values for *α* and *β*); 3) slow propagation of activity between the thalamus and cortex (high values for *t*_0_).

We employed a corticothalamic mean-field model that describes the mean membrane potential of four connected neuronal populations. Details of the model and the parameter estimation method can be found in the methods section. In brief, the model includes a cortical excitatory and cortical inhibitory population and a thalamic relay and reticular population (Figure 1C). The postsynaptic membrane potential of a population modulates as a consequence of synaptic input mediated by the firing activity of presynaptic populations. The effect of the presynaptic input on the postsynaptic membrane potential depends on the mean number of synapses between the presynaptic *b* and postsynaptic population *a* (modelled as synaptic strength *v*_*ab*_) and on the closing and opening rate of synaptic channels characterised by the synaptic decay and rise constants (*α* and *β*). The average postsynaptic membrane potential is transformed at the cell bodies in a population, giving rise to the firing activity. This process results in propagation of activity in a closed loop between thalamic and cortical populations, where firing rate propagated between the thalamus and the cortex is delayed by *t*_0_. Using *v*_*ab*_, we can disentangle this corticothalamic system to a purely cortical loop or gain *X*, a corticothalamic gain *Y* and a intrathalamic gain *Z*. The cortical gain is defined in terms of the ratio of cortical excitatory versus cortical inhibitory synaptic strengths.

Using a Markov Chain Monte Carlo random walk method^19–21^, we estimated the temporal evolution of these six biophysical parameters *X,Y, Z, α, β*, and *t*_0_ from consecutive EEG power spectra from every subject over the course of 48 hours. Figure 1D shows the goodness-of-fit in terms of *χ*^2^ for patients with a good and a poor outcome respectively. No significant difference was observed in model prediction accuracy (Mann-Whitney U *p >* 0.05).

For every parameter, the mean and standard error across subjects is depicted in Figure 1D, alongside the distribution of values from healthy controls (blue violin plots). Differences between groups across time points were assessed by non-parametric permutation testing^22^. For the cortical gain *X*, we see a separation between the means and their standard error for both groups in the initial phase (*t <* 24 hours after cardiac arrest), followed by an overlap between the means and their standard error for later phases (*t >* 24 hours after cardiac arrest). Hence, there is a higher cortical excitation inhibition ratio in patients with a poor outcome in the initial phase of *t <* 24 after cardiac arrest. Permutation testing across all time points, however, showed no significant difference between groups (p*>*0.05). For the corticothalamic gain *Y* there is no overlap between groups across all time points, resulting in a significant difference (p = 0.03). *Y* values for patients with a good outcome overlapped with the distribution of *Y* values from healthy controls, which was not the case for patients with a poor outcome. For patients with a poor outcome, *Y* is negative in contrast to positive values of *Y* for patients with a good outcome. This reflects dominance of inhibition in the corticothalamic loop in patients with a poor outcome; positive values correspond to dominance of excitation in the corticothalamic loop. For the intrathalamic gain *Z* there was a upward trend for patients with a good outcome and a downward trend for patients with a poor outcome. However, there was no significant difference between groups (p*>*0.05). For both patients with a good and poor outcome, the values of *Z* overlapped with the distribution of *Z* from healthy controls.

For the synaptic decay time *α* we observed a significant difference between patients with a good and a poor outcome (*p <* 0.001). Patients with a good outcome had lower values of *α*, with the tendency to reach the range of values of *α* of healthy control subjects at an earlier stage than patients with a poor outcome. For the synaptic rise time *β* there was a significant difference between groups (*p <* 0.001), with a steeper negative curve for patients with a good outcome and a tendency towards values of *β* from healthy controls. Lower values for *α* and *β* correspond to a synaptic response with smaller width, hence the ability to generate faster rhythms. The parameter *t*_0_ is the only parameter indicative of axonal integrity and captures the conduction velocity between the thalamus and the cortex. There was a significant difference in *t*_0_ values between groups (*p <* 0.001). Patients with a good outcome had overall shorter delays than patients with poor outcome. Around t = 48 h, patients with a good outcome had values of *t*_0_ that strongly overlapped with values of *t*_0_ from healthy control subjects.

## Discussion

We used a biophysical mean-field model to study the pathophysiology of EEG abnormalities in patients with a postanoxic encephalopathy, and show that poor outcome is associated with an initial high cortical excitation-inhibition ratio, relative inhibition in the corticothalamic loop, overall slow recovery of time scales of synaptic responses and longer delays in propagation of activity between the thalamus and cortex. Patients with a good outcome showed excitation in the corticothalamic loop, faster recovery of time scales of synaptic responses and faster propagation of activity between the cortex and thalamus with values that were essentially similar to those of healthy control subjects. Our findings show that poor outcome results from failure in synaptic recovery as well as from impaired axonal propagation (difference in *t*_0_ between groups).

In patients with a poor outcome, there was a higher cortical excitation-inhibition ratio in the first 24 hours after cardiac arrest compared to patients with a good outcome. This difference normalised after 24 hours. The higher cortical excitation-inhibition ratio is supported by clinical observations such as increased risk of electrographic or clinical seizures in this patient population^23, 24^, or high incidence of generalised periodic discharges, which may result from increased cortical excitability^6, 25^.

Our model suggests persistent corticothalamic synaptic failure in patients with a poor recovery. This finding is in line with animal work and postmortem data showing selective damage of reticular neurons in the thalamus after cardiac arrest^26, 27^, leading to excessive activity in the thalamus. Since efficient communication between the thalamus and the cortex depends on phasic inhibition or cyclic suppression in the thalamus^28^, allowing short temporal windows of sensitivity to synaptic input, tonic activity in the thalamus could disrupt this loop and hence could relate to decreased corticothalamic synaptic strength. Other post-mortem work also suggest that patients with thalamic damage had EEG abnormalities associated with poor outcome^9^. Moreover, integrity of corticothalamic synapses is associated with the ability to generate alpha oscillations^29, 30^ and usually emerge from the model if *Y >* 0^29^. This was in line with our empirical findings, as patients with a good outcome in general had *Y >* 0. Patients with a poor outcome had negative values for *Y*, which is usually associated with slower delta and theta activity^29^. We also show that the integrity of corticothalamic synaptic connections are more important to distinguish good from poor outcome than intra-cortical interactions. This in agreement with the so-called mesocircuit hypothesis^31, 32^. This hypothesis postulates that recovery of consciousness after severe brain injury (both postanoxic encephalopathy and traumatic brain injury) strongly depends on recovery of excitatory activity in a thalamo-cortical circuit.

Failure of synaptic transmission is a well-known consequence of cerebral hypoxia or ischemia^14, 33^. Here we report longer synaptic rise and decay times for all synapses in patients with a poor outcome. These parameters correspond to opening and closing times of ligand-gated channels in the synapse and can be translated to a synaptic impulse response function^34^. The opening and closing times of ligand-gated channels strongly relate to the frequency response of post-synaptic membranes and longer synaptic rise and decay times correspond to the inability to generate faster rhythms (alpha and beta) in patients with a poor outcome^35^. Results agree with previous animal and in-vitro work showing longer postsynaptic potentials (PSP) after hypoxia^36, 37^. This is probably the result of long-term potentiation (LTP) mediated by upregulation of NMDA-receptors due to excess of extracellular glutamate after hypoxia^38^. We show that opening times of ligand-gated synaptic channels are shorter and normalise faster in patients with a good outcome compared to poor outcome, but still differs from values from healthy controls. Hence, further recovery of the EEG may depend on reversing LTP^14^.

Propagation of activity between the cortex and thalamus was severely delayed for patients with a poor outcome. Longer delays in the corticothalamic loop generally result in slower theta and delta rhythms^35^, consistent with our empirical findings in this group with a poor outcome. For patients with a good outcome, there was evolution towards a conduction delay comparable to that of healthy control subjects. Previous work showed that mild hypoxia induces isolated synaptic failure with intact heights and shapes of the action potential, suggesting intact axonal membrane functioning^33^. Other in-vitro work has shown that oligodendrocytes are vulnerable to hypoxia, resulting in defects in myelination due to hypoxia^39, 40^. Animal work has also shown that anoxia can lead to detachment of perinodal oligodendrocyte-axon loops, and thus cause axonal injury^41^. Our finding of longer conduction delays between the cortex and thalamus in patients with a poor outcome is suggestive of myelination defects rather than axonal damage. This is in line with findings from diffusion tensor imaging studies showing increased radial diffusivity in patients after cardiac arrest, suggestive of myelination defects^42^.

Some limitations apply to our work. First, synaptic rise and decay times were agreggated for all populations (excitatory, inhibitory, and thalamic). This simplification is probably not justified by the underlying neurophysiology, but was required to reduce the number of estimated parameters. Second, we ignored spatial effects. Input to our model were power spectra averaged across all electrodes. While it may be more realistic to estimate sensor specific parameters, the EEG after cardiac arrest does not show high regional specificity but rather homogeneous activity over different electrodes. Furthermore, this would have resulted in an excess of parameters. Third, our model cannot disentangle presynaptic and postsynaptic effects. Last, in order to estimate underlying neurophysiological parameters, we did not employ the full nonlinear model, but a linearised version of the model. This is probably not critical as previous work suggests that many empirical phenomenon can be captured using the linearised version of the mean-field model^35, 43^.

In conclusion, we have provided insight into potential pathophysiological mechanisms of EEG abnormalities in postanoxic encephalopathy. We show that these EEG abnormalities are mostly reflective of synaptic failure, though not limited to isolated synaptic failure as poor outcome was also accompanied with longer axonal conduction delays between the cortex and thalamus, probably reflecting myelination defects. Preservation of thalamocortical synaptic connections, propagation of neuronal activity between the cortex and thalamus and normalisation of synaptic responses appear crucial for evolution of the EEG associated with good outcome in patients with postanoxic encephalopathy. This framework and these findings pave the way to detect synaptic failure in individual patients.

## Methods

### Study population

We selected EEGs from 50 comatose survivors of cardiac arrest with poor outcome and 50 patients with good outcome from our previously published dataset^3, 6^. Selection was based on three criteria: a Cerebral Performance Score (CPC) of either 1 (poor outcome) or 5 (good outcome) at six months after cardiac arrest to maximise contrast between groups; CPC of 1 only as a result of postanoxic coma (and not for example as a result of multiple organ or systemic failure); and an evolution towards a continuous or discontinuous EEG at 48 hours post-cardiac arrest. This last criterium is to ensure that we avoid mixing up different EEG abnormalities and corresponding pathways in patients with poor outcome. For example, the pathophysiological underpinnings burst-suppression with identical bursts may be different from the pathophysiological underpinnings of a continuous EEG with GPDs. Hence, EEGs with the following abnormalities were excluded during our selection: GPDs on an isoelectric background, burst-suppression as final stage, rhythmic activity (*>* 2.5 Hz) or periodic activity (0.5 − 2.5 Hz), a suppressed EEG. To obtain a reference for all estimated biophysical parameters, we used eyes-closed EEG data from 100 healthy controls.

### EEG preprocessing

Continuous EEG recordings were used from patients with cardiac arrest admitted to the ICU. Nineteen electrodes (either silver/silver chloride cup or subdermal wire) were placed according to the 10–20 International System. A Neurocenter EEG system with Refa or SAGA amplifiers (TMSi, Netherlands) was used, recording at a sample frequency of 256 Hz. EEG data until 48 hours after cardiac arrest was used for further analysis using the longitudinal bipolar montage. These EEG data were further preprocessed using a zero-phase sixth-order Butterworth bandpass filter of 0.5-40 Hz. We used a semi-automated algorithm to detect and remove artifacts within windows of 10 seconds in the common average. Artifacts included empty channels, channels with large peaks or noise (amplitude ≥ 200 µV or ≤ − 200 µV and variance ≥ 1400 µV^2^ or ≤ 1 µV^2^), or muscle artifacts. In addition, we used independent component analysis to detect and remove the ECG artefact after visual inspection of individual components^44^. After pre-processing, we computed a power spectrum for every hour and every channel and averaged across channels. Power spectra were computed using Welch’s method with windows of 10 seconds with 5 seconds overlap.

### Corticothalamic mean-field model

We employed a corticothalamic mean-field model^35, 45, 46^, which describes the aggregate activity of a neuronal population in terms of their firing activity *φ*_*a*_ and mean membrane potential *V*_*a*_ with *a* ∈ {*e, i, r, s*}. The corticothalamic mean-field model encompasses two cortical populations (excitatory *e*, inhibitory *i*) and two thalamic populations (relay *s*, reticular *r*). The membrane potential of a population fluctuates *V*_*a*_(*t*) as a result of the incoming firing rate *φ*_*a*_(*t*) from other population and/or itself according to

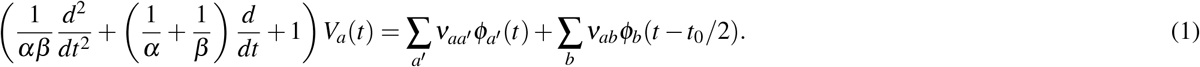

The effect of the presynaptic input on the postsynaptic membrane potential depends on the mean number of synapses between the presynaptic *b* and postsynaptic population *a* (modelled as synaptic strength *v*_*ab*_) and on the closing and opening rate of synaptic channels characterised by the synaptic decay and rise constants (*α* and *β*). The average postsynaptic membrane potential is transformed at the cell bodies in a population, giving rise to the firing activity *Q*_*a*_(*t*).

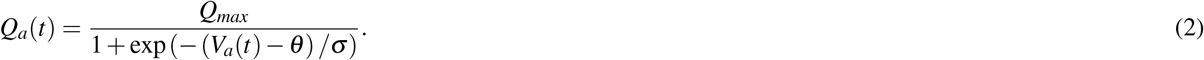

This process results in propagation of activity in a closed loop between thalamic and cortical populations, where firing rate propagated between the thalamus and the cortex is delayed by *t*_0_. The firing activity *Q*_*a*_(*t*) is further temporally damped using the following expression

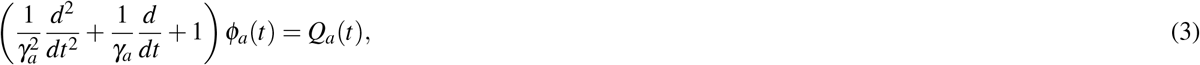

with *γ*_*a*_ being the temporal damping rate, based on *γ* = *v*_*a*_*/r*_*a*_, where *v*_*a*_ is the propagation velocity and *r*_*a*_ is the mean range of axons. For inhibitory, relay and reticular populations, *γ*_*a*_ *≈* ∞, hence *φ*_*a*_(*t*) = *Q*_*a*_(*t*).

### Parameter estimation of model parameters

#### Model power spectrum

Parameter estimation for nonlinear models remains challenging. For example, error of estimated parameters near critical points or bifurcations can have severe effect on the expected behaviour of the model. Equations (1-3) describe the full nonlinear model, which are first transformed to a linear model using linearisation around a stable fixed point. Linearisation is achieved by expressing the sigmoid function (Equation 2) that transforms *V*_*a*_(*t*) into *Q*_*a*_(*t*) as Taylor expansion and retaining only the term containing the first derivative *ρ*_*a*_ evaluated at the fixed point. Details can be found in^19^. Using the derivative *ρ*_*a*_, we can express the synaptic strengths as gain parameters in the linear regime *G*_*ab*_ = *ρ*_*a*_*ν*_*ab*_. These gain parameters can be transformed to three gain parameters describing the cortical *X*, corticothalamic *Y* and intrathalamic loop gains

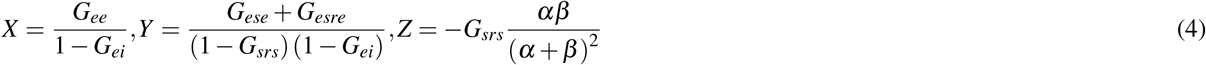

The cortical gain is defined in terms of the ratio of cortical excitatory versus cortical inhibitory synaptic strengths. Following this, the linear system in time domain can be rewritten in Fourier domain from which we can express an analytical expression for the power spectrum *P*(*w*) as a function of frequency *ω*

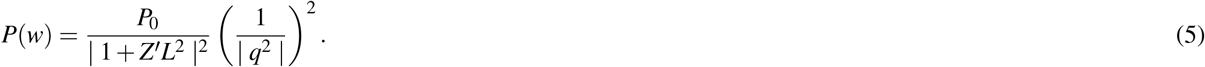

*P*_0_ is a normalisation constant^46^ and *q* follows from the dispersion relation described in^19^

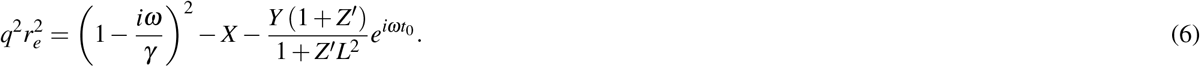

*Z*^*′*^ follows from a transformation of *Z*

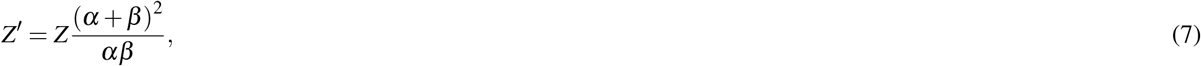

and *L* follows from the transformation of the second order differential operator describing the synaptic response (Equation 1) in Fourier domain, which can be interpreted as a low-pass filter depending on the synaptic parameters *α* and *β*

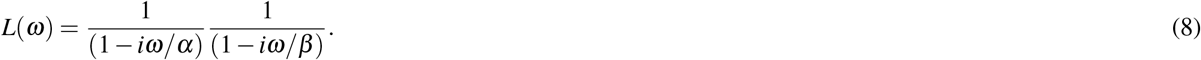

Activity in pericranial muscles results in electromyogram (EMG) artefacts on the EEG, for which we need to account, hence, *P*_*total*_ (*ω*) = *P* (*ω*) + *P*_*EMG*_ (*ω*).

#### Fitting model parameters

The parameter set **x** = [*X,Y, Z, α, β, t*_0_, *P*_*EMG*_] is estimated from EEG data by minimising the error between the experimentally obtained power spectrum *P*_*exp*_ and model power spectrum *P*_*total*_ (**x**) expressed as

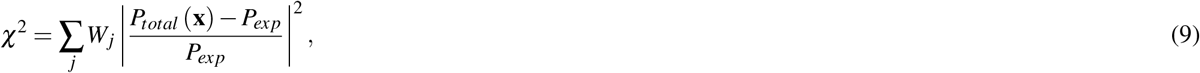

where *j* indexes the frequency bins. The weights *W*_*j*_ ensure equal weighting for every frequency decade and is proportional to 1*/ f*. As parameter space is very large, we restrict parameter values to neurophysiologically plausible values (see^19^ for values). The *χ*^2^ statistic is further transformed into a likelihood function

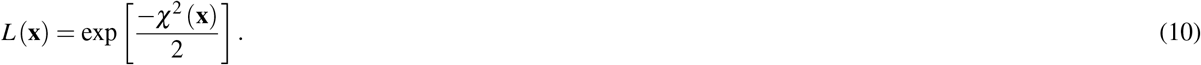

Hence, minimising the error translates into maximising this likelihood function. Now, the Metropolis-Hastings algorithm is used to generate a probability distribution for each parameter using a Markov Chain random walk^20, 21^. Details of the algorithm can be found in^19^. For every subject, we run the Metropolis-Hastings algorithm to obtain model parameters for individual power spectra. The random walk is initialised by parameters obtained from a large database of healthy control subjects^43^. This initialisation will generally not affect the final output, but it affects the time of convergence. For every subsequent step in the random walk, the likelihood for this step is computed using Equation 10. A new random proposed set is generated. The likelihood of this new set of parameters is again computed using Equation 10. If these new parameters have a higher probability, this step is accepted and used to sample the probability distribution. Otherwise a random number is drawn from a uniform distribution. If this random number is smaller than the ratio of the probability of the new parameters to the old parameters, accept the step for sampling the probability distribution. If this random number is bigger than the ratio of the probability of the new parameters to the old parameters, then this step is not accepted. This procedure is repeated many times until there is no iterative change in the sampled probability distribution. Fitting a sequence of spectra at successive times would require to track temporal changes in parameter values. If we assume that the power spectrum does not change drastically for consecutive time points *t*_*i*_ and *t*_*i*+1_, then we can use Bayes’s theorem to inform our fit for *t*_*i*+1_ using estimated parameters from *t*_*i*_ as prior information. Code for the parameter estimation method can be found on https://github.com/BrainDynamicsUSYD/braintrak, which includes explanation of the method on the wiki page https://github.com/BrainDynamicsUSYD/braintrak/wiki.

#### Statistics

We used non-parametric permutation testing to test significance between groups of patients with a good and poor outcome^22^. We tested whether a parameter across time points was significantly different between groups. We did not test for significance between groups for every time point separately to avoid a multiple testing problem. For every time dependent parameter *x*_*s jg*_(*t*), with parameter index *j*, we computed the mean across subjects *s* for every group *g* denoted 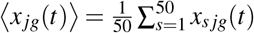. We subsequently used the sum of the squared difference between group *g* (good outcome) and group *g*^*′*^ (poor outcome) as test statistic *T*_*j*_ = ∑ (*⟨x* _*jg*_(*t*) *⟩ −* ⟨ *x* _*jg*_^*′*^ (*t*) ⟩)^2^. Following this, we randomly permuted group membership 10 000 times and computed the test statistic for every realisation to generate a null-distribution. The genuine value of the test statistic *T*_*j*_ was subsequently compared to the null distribution and was considered to be significant if this would lie in the two 2.5% tail-ends of the null-distribution (*p <* 0.05). We performed in total six statistical tests (parameters *X,Y, Z, α, β, t*_0_). All statistics were performed in MATLAB R2021a.

## Acknowledgements

We would like to acknowledge Carin J. Eertman-Meyer for data acquisition. We would like to thank Barry Ruijter, MD PhD for providing pre-processed, relative artefact free EEG data.

## References

1. Sandroni, C. et al. Prediction of poor neurological outcome in comatose survivors of cardiac arrest: a systematic review. Intensive care medicine 46, 1803–1851 (2020).

2. Hofmeijer, J. et al. Early eeg contributes to multimodal outcome prediction of postanoxic coma. Neurology 85, 137–143 (2015).

3. Tjepkema-Cloostermans, M. C. et al. Electroencephalogram predicts outcome in patients with postanoxic coma during mild therapeutic hypothermia. Critical care medicine 43, 159–167 (2015).

4. Jørgensen, E. & Holm, S. The natural course of neurological recovery following cardiopulmonary resuscitation. Resuscitation 36, 111–122 (1998).

5. Sandroni, C. et al. Prediction of good neurological outcome in comatose survivors of cardiac arrest: a systematic review. Intensive care medicine 1–25 (2022).

6. Ruijter, B. J., van Putten, M. J. & Hofmeijer, J. Generalized epileptiform discharges in postanoxic encephalopathy: quantitative characterization in relation to outcome. Epilepsia 56, 1845–1854 (2015).

7. Ruijter, B. J. et al. Early electroencephalography for outcome prediction of postanoxic coma: a prospective cohort study. Annals neurology 86, 203–214 (2019).

8. Zandt, B.-J., ten Haken, B., van Dijk, J. G. & van Putten, M. J. Neural dynamics during anoxia and the “wave of death”. PLoS One 6, e22127 (2011).

9. van Putten, M. J. et al. Postmortem histopathology of electroencephalography and evoked potentials in postanoxic coma. Resuscitation 134, 26–32 (2019).

10. Dijkstra, K., Hofmeijer, J., van Gils, S. A. & van Putten, M. J. A biophysical model for cytotoxic cell swelling. J. neuroscience 36, 11881–11890 (2016).

11. Nutma, S., Le Feber, J. & Hofmeijer, J. Neuroprotective treatment of postanoxic encephalopathy: a review of clinical evidence. Front. neurology 12, 614698 (2021).

12. Hofmeijer, J. & van Putten, M. J. Ischemic cerebral damage: an appraisal of synaptic failure. Stroke 43, 607–615 (2012).

13. van Putten, M. J. & Hofmeijer, J. Eeg monitoring in cerebral ischemia: basic concepts and clinical applications. J. clinical neurophysiology 33, 203–210 (2016).

14. Ruijter, B. J., Hofmeijer, J., Meijer, H. G. E. & van Putten, M. J. A. M. Synaptic damage underlies eeg abnormalities in postanoxic encephalopathy: A computational study. Clin. neurophysiology 128, 1682–1695 (2017).

15. Coombes, S. Large-scale neural dynamics: simple and complex. NeuroImage 52, 731–739 (2010).

16. Deco, G., Jirsa, V. K., Robinson, P. A., Breakspear, M. & Friston, K. The dynamic brain: from spiking neurons to neural masses and cortical fields. PLoS computational biology 4, e1000092 (2008).

17. Tjepkema-Cloostermans, M. C., Hindriks, R., Hofmeijer, J. & van Putten, M. J. Generalized periodic discharges after acute cerebral ischemia: reflection of selective synaptic failure? Clin. neurophysiology 125, 255–262 (2014).

18. Victor, J. D., Drover, J. D., Conte, M. M. & Schiff, N. D. Mean-field modeling of thalamocortical dynamics and a model-driven approach to eeg analysis. Proc. Natl. Acad. Sci. 108, 15631–15638 (2011).

19. Abeysuriya, R. & Robinson, P. Real-time automated eeg tracking of brain states using neural field theory. J. neuroscience methods 258, 28–45 (2016).

20. Metropolis, N., Rosenbluth, A. W., Rosenbluth, M. N., Teller, A. H. & Teller, E. Equation of state calculations by fast computing machines. The journal chemical physics 21, 1087–1092 (1953).

21. Rosenthal, J. S. et al. Optimal proposal distributions and adaptive mcmc. Handb. Markov Chain Monte Carlo 4 (2011).

22. Nichols, T. E. & Holmes, A. P. Nonparametric permutation tests for functional neuroimaging: a primer with examples. Hum. brain mapping 15, 1–25 (2002).

23. Rossetti, A. O. et al. Status epilepticus: an independent outcome predictor after cerebral anoxia. Neurology 69, 255–260 (2007).

24. Ruijter, B. J. et al. Treating rhythmic and periodic eeg patterns in comatose survivors of cardiac arrest. New Engl. J. Medicine 386, 724–734 (2022).

25. Barbella, G. et al. Prediction of regaining consciousness despite an early epileptiform eeg after cardiac arrest. Neurology 94, e1675–e1683 (2020).

26. Shoykhet, M. et al. Thalamocortical dysfunction and thalamic injury after asphyxial cardiac arrest in developing rats. J. Neurosci. 32, 4972–4981 (2012).

27. Ross, D. T. & Graham, D. I. Selective loss and selective sparing of neurons in the thalamic reticular nucleus following human cardiac arrest. J. Cereb. Blood Flow & Metab. 13, 558–567 (1993).

28. Lőrincz, M. L., Kékesi, K. A., Juhász, G., Crunelli, V. & Hughes, S. W. Temporal framing of thalamic relay-mode firing by phasic inhibition during the alpha rhythm. Neuron 63, 683–696 (2009).

29. Roberts, J. & Robinson, P. Corticothalamic dynamics: structure of parameter space, spectra, instabilities, and reduced model. Phys. Rev. E 85, 011910 (2012).

30. Hindriks, R. & van Putten, M. J. Thalamo-cortical mechanisms underlying changes in amplitude and frequency of human alpha oscillations. Neuroimage 70, 150–163 (2013).

31. Schiff, N. D. Recovery of consciousness after brain injury: a mesocircuit hypothesis. Trends neurosciences 33, 1–9 (2010).

32. Edlow, B. L., Claassen, J., Schiff, N. D. & Greer, D. M. Recovery from disorders of consciousness: mechanisms, prognosis and emerging therapies. Nat. Rev. Neurol. 17, 135–156 (2021).

33. Hofmeijer, J., Mulder, A. T., Farinha, A. C., van Putten, M. J. & le Feber, J. Mild hypoxia affects synaptic connectivity in cultured neuronal networks. Brain research 1557, 180–189 (2014).

34. Rennie, C. J., Robinson, P. A. & Wright, J. J. Unified neurophysical model of eeg spectra and evoked potentials. Biol. cybernetics 86, 457–471 (2002).

35. Robinson, P. A. et al. Prediction of electroencephalographic spectra from neurophysiology. Phys. Rev. E 63, 021903 (2001).

36. Urban, L., Neill, K., Crain, B., Nadler, J. & Somjen, G. Postischemic synaptic physiology in area ca1 of the gerbil hippocampus studied in vitro. J. Neurosci. 9, 3966–3975 (1989).

37. Miyazaki, S. et al. Post-ischemic potentiation of schaffer collateral/ca1 pyramidal cell responses of the rat hippocampus in vivo: involvement of n-methyl-d-aspartate receptors. Brain research 611, 155–159 (1993).

38. Szatkowski, M. & Attwell, D. Triggering and execution of neuronal death in brain ischaemia: two phases of glutamate release by different mechanisms. Trends neurosciences 17, 359–365 (1994).

39. Dewar, D., Underhill, S. M. & Goldberg, M. P. Oligodendrocytes and ischemic brain injury. J. Cereb. Blood Flow & Metab. 23, 263–274 (2003).

40. Waxman, S. G., Black, J. A., Ransom, B. R. & Stys, P. K. Anoxic injury of rat optic nerve: ultrastructural evidence for coupling between na+ influx and ca2+-mediated injury in myelinated cns axons. Brain research 644, 197–204 (1994).

41. Lyons, S. A. & Kettenmann, H. Oligodendrocytes and microglia are selectively vulnerable to combined hypoxia and hypoglycemia injury in vitro. J. Cereb. Blood Flow & Metab. 18, 521–530 (1998).

42. Laitio, R. et al. Effect of inhaled xenon on cerebral white matter damage in comatose survivors of out-of-hospital cardiac arrest: a randomized clinical trial. Jama 315, 1120–1128 (2016).

43. Abeysuriya, R., Rennie, C. & Robinson, P. Physiologically based arousal state estimation and dynamics. J. Neurosci. Methods 253, 55–69 (2015).

44. Hyvarinen, A. Fast and robust fixed-point algorithms for independent component analysis. IEEE transactions on Neural Networks 10, 626–634 (1999).

45. Freyer, F. et al. Biophysical mechanisms of multistability in resting-state cortical rhythms. J. Neurosci. 31, 6353–6361 (2011).

46. Robinson, P., Rennie, C. & Rowe, D. Dynamics of large-scale brain activity in normal arousal states and epileptic seizures. Phys. Rev. E 65, 041924 (2002).

